# Purine-responsive expression of the *Leishmania donovani* NT3 purine nucleobase transporter is mediated by a conserved RNA stem-loop

**DOI:** 10.1101/2020.01.26.920504

**Authors:** M. Haley Licon, Phillip A. Yates

**Affiliations:** Department of Molecular Microbiology & Immunology, Oregon Health & Science University, Portland OR 97239; Department of Chemical Physiology & Biochemistry, Oregon Health & Science University, Portland OR 97239

**Keywords:** Leishmania, post-transcriptional regulation, purine, mRNA, parasitology, stress response, transporter, adaptation, starvation

## Abstract

The ability to modulate gene expression in response to changes in the host environment is essential for survival of the kinetoplastid parasite *Leishmania*. Unlike most eukaryotes, gene expression in kinetoplastids is predominately regulated post-transcriptionally. Consequently, RNA-binding proteins (RBPs) and mRNA-encoded sequence elements serve as primary determinants of gene regulation in these organisms; however, few have been ascribed roles in specific stress-response pathways. *Leishmania* lack the capacity for *de novo* purine synthesis and must scavenge these essential nutrients from the host. *Leishmania* have evolved a robust stress response to withstand sustained periods of purine scarcity during their lifecycle. The purine nucleobase transporter, LdNT3, is among the most substantially upregulated proteins in purine-starved *L. donovani.* Here we report that the post-translational stability of the LdNT3 protein is unchanged in response to purine starvation. Instead, LdNT3 upregulation is primarily mediated by a 33 nucleotide (nt) sequence in the *LdNT3* mRNA 3’-untranslated region that is predicted to adopt a stem-loop structure. While this sequence is highly conserved within the mRNAs of orthologous transporters in multiple kinetoplastid species, putative stem-loops from *L. donovani* and *Trypanosoma brucei* nucleobase transporter mRNAs are not functionally interchangeable for purine-responsive regulation. Through mutational analysis of the element, we demonstrate that species specificity is attributable to just three variant bases within the predicted loop. Finally, we provide evidence that the abundance of the *trans-*acting factor that binds the *LdNT3* stem-loop *in vivo* is substantially higher than required for regulation of LdNT3 alone, implying a potential role in regulating other purine-responsive genes.

---

Kinetoplastid parasites of the genus *Leishmania* are the etiological agents of leishmaniasis, a suite of debilitating and often fatal diseases that affect roughly 12 million people worldwide (1). Over the course of their lifecycles, *Leishmania* exist as both extracellular promastigotes in the ambient midgut of a sandfly vector and as intracellular amastigotes in the phagolysosomes of mammalian macrophages. As these compartments differ substantially in pH, temperature, and nutrient availability, *Leishmania* must undergo dramatic metabolic and physiological transformations to adapt to life in their respective hosts (2,3). Such environmental responses are so intertwined with the leishmanial lifecycle that several stressors function as cues for parasite differentiation, triggering the transition from one developmental stage to the next (4-8). Despite their importance to parasite biology, relatively little is known of the molecular mechanisms that allow *Leishmania* to respond to fluctuations in the extracellular milieu.

*Leishmania* and other kinetoplastid protozoa separated from the eukaryotic lineage early in evolutionary history and consequently exhibit several unique biological features, including their means of gene expression. Deviating from the one-promotor-to-one-gene canon of higher eukaryotes, the genomes of kinetoplastid parasites are organized as large unidirectional transcription units that can comprise tens to hundreds of tandemly arranged genes and span up to 100 kb in length (9). Transcription by RNA polymerase II results in the production of polycistronic primary transcripts, which mature into single-gene mRNAs through 3’ polyadenylation and *trans*-splicing of a capped exon (the spliced leader) to the 5’ end of each coding sequence (CDS) (reviewed in 10). As a result, *Leishmania* rely almost exclusively on post-transcriptional control points like mRNA stability and localization, translational efficiency, and protein half-life to respond to exogenous stimuli. As in higher eukaryotes, transcript abundance and translation in the kinetoplastids are modulated by the interactions of *trans*-acting RBPs and *cis-*acting regulatory elements encoded in mRNA UTRs or the CDS itself. In recent decades, much effort has been devoted to the study of such elements in these organisms; however, only a handful of RBPs and even fewer discrete mRNA elements have been identified as both necessary and sufficient for gene regulation in the context of specific stress response pathways (11-13).

*Leishmania* are unable to synthesize purines *de novo* and must scavenge these essential nutrients from the host (14, 15). The availability of salvageable purines likely fluctuates throughout the parasite lifecycle and *Leishmania* have evolved to withstand periods of extreme purine scarcity. Indeed, this particular type of nutrient stress may be an important regulator of parasite infectivity. Adenosine supplementation negatively impacts the efficiency of *in vivo* metacyclogenesis, the process wherein poorly infective procyclic promastigotes in the sandfly midgut differentiate into highly infective metacyclic promastigotes, primed for survival within a vertebrate host (8). A robust and reproducible purine stress response is easily induced *in vitro* by omission of purines from the growth medium, and we showed that purine-starved *Leishmania donovani* promastigotes enter a reversible quiescent-like state in which they can persist for over three months (14, 16, 17). Whole proteome comparisons of purine-starved and −replete *L. donovani* promastigotes conducted over a 6-to 48-hour window revealed a temporally-controlled remodeling of the cellular proteome. While early changes centered on increasing purine uptake, later responses reflected a larger restructuring of cellular metabolism to reduce energy expenditure and enhance general stress tolerance. The global transcriptomes of these cells were also significantly different; however, changes in mRNA abundance often tracked poorly with those manifested at the protein level, implicating both translational and post-translational mechanisms (16). Together, these analyses demonstrated that purine-responsive cellular remodeling is complex and orchestrated at multiple levels of post-transcriptional regulation.

Though our earlier global analyses provided critical insight into *what* processes are important for adaptation to purine starvation, they uncovered little about *how* specific changes in protein abundance are established. In this report, we look deeper at the molecular mechanisms underlying the leishmanial purine stress response through the lens of a representative purine-responsive gene. The *L. donovani* purine nucleobase transporter 3 (LdNT3) is one of the earliest and most substantially upregulated proteins in purine-starved *L. donovani.* LdNT3 upregulation is mediated in part at the levels of mRNA abundance and translational efficiency (16, 18). Here we demonstrate that LdNT3 protein stability is not altered under purine stress, establishing the dominance of mRNA-level and translational control points in purine-responsive LdNT3 regulation. We identify a 33 nt predicted stem-loop sequence in the *LdNT3* 3’-UTR (referred to as the LdNT3 stem-loop) that represses expression when extracellular purines are abundant. Using a series of integrating luciferase reporter constructs, we show that the *LdNT3* stem-loop is sufficient to confer purine-responsive regulation in heterologous sequence contexts. We examine evolutionary conservation of the element and conduct a thorough mutational analysis to identify functionally important regions required for repressor activity in purine-replete *L. donovani*. Lastly, we show that the *LdNT3* stem-loop is sufficient to confer purine-responsiveness to a high-abundance transcript, suggesting that the cognate RBP responsible for binding this element *in vivo* is present within the cell in substantial excess of what is required to regulate *LdNT3* expression alone.

## RESULTS

### LdNT3 protein stability is not regulated in response to purine stress

To elucidate the molecular mechanisms that coordinate purine-responsive LdNT3 expression, we first determined the levels at which they operate. It is published that changes in both mRNA stability and translational efficiency contribute to LdNT3 upregulation under purine starvation (16, 18). However, our earlier studies ignored the potential contribution of post-translational stabilization. In our experience, epitope-tagged LdNT3 is refractory to direct detection by western blotting at the low levels expressed in purine-replete cultures. Therefore, we modified a dual-luciferase system established in our laboratory to indirectly measure changes in LdNT3 protein stability via enzymatic reporter assay.

In the simplest iteration of the dual-luciferase system (described at length in 18), the firefly luciferase gene (*Fluc*) is fused in-frame with a selectable drug resistance marker and integrated in place of one allelic copy of the gene of interest. Reporter integration fully replaces the CDS of the targeted gene while preserving the endogenous intergenic regions (IGRs), which contain the requisite signals for trans-splicing and polyadenylation. Importantly, as the 5’ and 3’ mRNA UTRs are derived from these up- and downstream IGRs, respectively, their contributions to regulation are reflected in luciferase activity. To normalize Fluc activity between replicates, *Renilla* luciferase (*Rluc*) is similarly integrated into the locus of a gene for which expression does not change under the conditions of the experiment. In probing the purine stress response, we used UMP synthase (*UMPS*) as an unresponsive control, since neither UMPS mRNA nor protein abundance are affected by purine starvation (16, 18).

To adapt this system for the study of post-translational stability, we generated cell lines in which the *LdNT3* CDS was fused via its N-terminus to a Fluc reporter and integrated into either its endogenous locus or that of a purine-unresponsive control, *LdNT4* (Figure 1A). The multicistronic constructs used for integration contained a blasticidin resistance gene (*BSD*) to facilitate mutant selection and a 2A peptide from the *Thosea asigna* virus (*2A*) that is co-translationally cleaved, liberating the BSD-2A polypeptide to minimize the size of appended tag on LdNT3 (19, 20). In this configuration, the post-translational fate of the reporter is coupled to that of the transporter while mRNA stability and translation are governed by the native *LdNT3* UTRs and/or CDS such that changes in luciferase activity reflect the cumulative effect of mechanisms operating at all post-transcriptional levels. In contrast, the purine nucleobase transporter 4 (LdNT4), while homologous to LdNT3, is not differentially regulated with respect to purine availability and Fluc expression from this locus consistently reflects an absence of purine sensitivity (16). Thus, expression of Fluc-LdNT3 integrated into this locus reflects only regulation conferred by elements contained within the *LdNT3* CDS itself or directly affecting stability of the protein. As anticipated, endogenously tagged Fluc-LdNT3 was significantly upregulated by 24 hours of purine starvation, consistent with previous experiments that implicated the UTRs in LdNT3 regulation. In cells expressing *Fluc-LdNT3* flanked by neutral *LdNT4* UTRs, however, luciferase activity was not affected by purine stress (Figure 1B). These data indicate that the *LdNT3* CDS does not encode additional *cis*-acting purine response elements nor does protein stability contribute to LdNT3 upregulation in purine-starved parasites. Thus, in the absence of post-translational control, transcript stability and translational efficiency serve as primary control points mediating purine-responsive changes in LdNT3 abundance.

**Figure 1.**
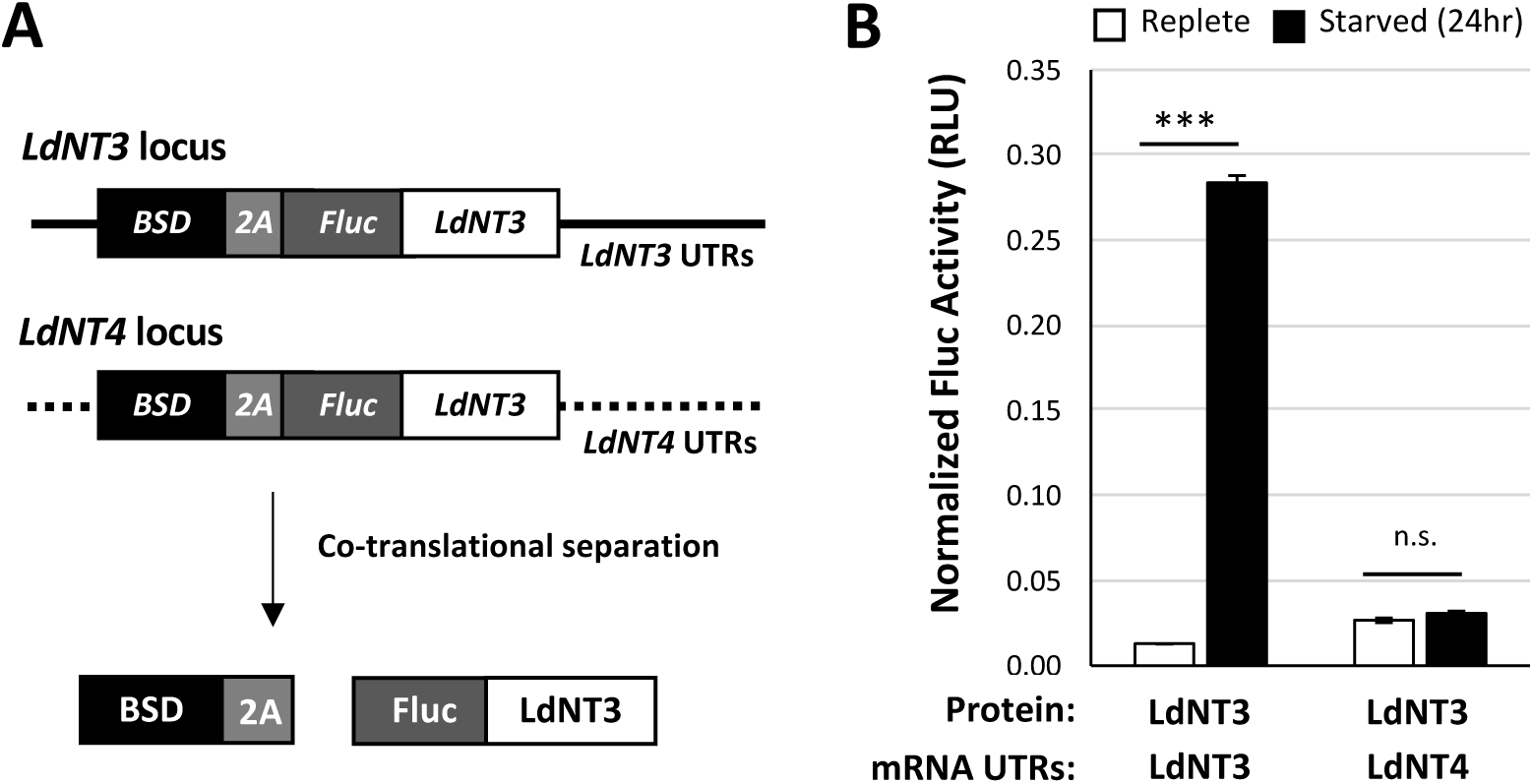
Changes in protein stability do not contribute to LdNT3 upregulation in purine-starved *Leishmania donovani.* A) Reporter constructs integrated into either the *LdNT4* or *LdNT3* locus to distinguish the relative contribution of protein stability from the cumulative effect of all post-transcriptional control points on LdNT3 upregulation. Solid and dashed lines indicate purine-responsive and -unresponsive mRNA UTRs, respectively. Not pictured: In this and all subsequent experiments, a *Renilla* luciferase-puromycin resistance gene fusion (*Rluc-PAC)* expressed from the *UMPS* locus serves as an internal normalization control. B) Luciferase activity from cell lines depicted in A, after 24 hours of culture in the presence (replete) or absence (starved) of purines. Figure shows the mean and standard deviation of experiments performed in biological and technical duplicate. Asterisks (*) indicate significance: single-factor ANOVA calculated with Excel Descriptive Statistics Toolpak; n.s., not significant, *P* ≥ 0.05; \*\*\**P* ≤ 0.001.

### A 33 nt stem loop in the *LdNT3* mRNA 3’-UTR represses expression under purine-replete conditions

The *LdNT3* 5’- and 3’-UTRs are together sufficient to confer purine-responsiveness to a reporter (16, 18), strongly implicating the presence of *cis*-acting regulatory sites within one or both of these regions. The 3’-UTR of the orthologous *Trypanosoma brucei* purine nucleobase transporter, *TbNT8.1*, encodes a predicted stem-loop that both is necessary and sufficient to repress TbNT8.1 expression when extracellular purines are abundant (13). Based on homology to this region, we identified a similar 33 nt predicted stem-loop in the *LdNT3* 3’-UTR, approximately 2.73 kb downstream of the stop codon. (Figures 2A and 2B). While the exact secondary structure of this element *in vivo* has not been formally demonstrated, we will refer to these sequences as stem-loops throughout for convenience. Though absent from the UTRs of other purine-responsive genes in *L. donovani*^*1*^, this element was identified in the UTRs of orthologous purine transporters from a variety of kinetoplastids, suggesting a strong evolutionary pressure for conservation (Figure 2A; Figure S1). To test whether this region also confers purine-responsiveness in *L. donovani*, we generated cell lines in which a *Fluc-BSD* transgene was expressed from the endogenous *LdNT3* locus under the control of either wildtype UTRs or a modified 3’-UTR lacking the putative stem-loop (Figure 2B). Parasites expressing *Fluc-BSD* flanked by wildtype UTRs demonstrated an approximate 12-fold increase in luciferase activity in response to purine stress. The magnitude of this effect was diminished by nearly ninety percent in stem-loop deletion mutants, wherein Fluc activity was increased only ∼1.4-fold by starvation. Specifically, deletion of the stem-loop resulted in a ∼11-fold increase in basal Fluc expression under purine-replete conditions, consistent with the sequence functioning as a negative regulator (Figure 2C). Together, these data implicate the repressive *LdNT3* stem-loop is a major regulator of purine-responsive LdNT3 expression.

**Figure 2.**
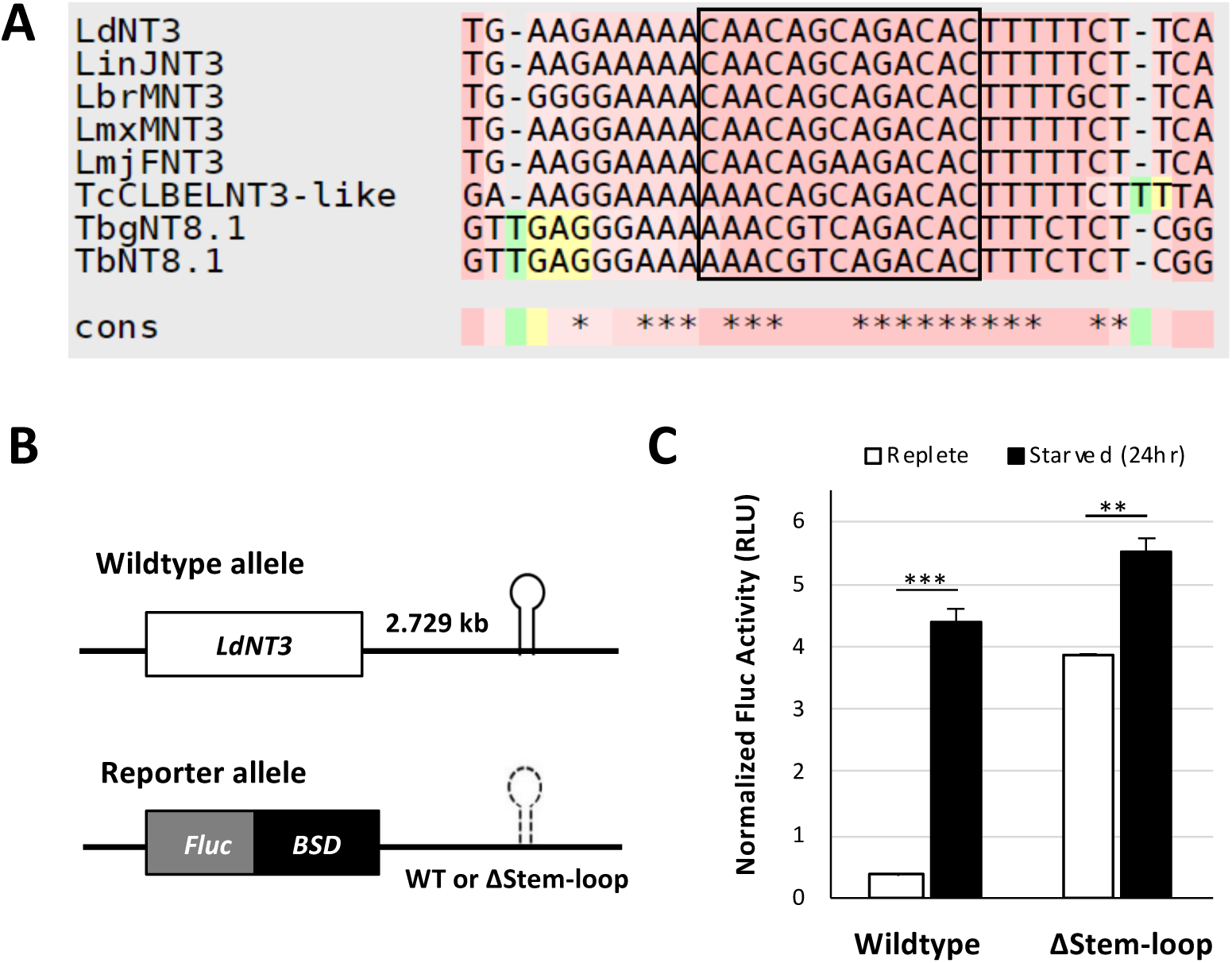
A 33 nt stem-loop in the *LdNT3* mRNA 3’-UTR represses LdNT3 expression under purine-replete conditions. A) Multiple sequence alignment of putative purine-response elements from *Leishmania donovani* (LdNT3; LdBPK_131110.1), *L. infantum* (LinJNT3; LINF_130017100), *L. braziliensis* (LbrMNT3; LbrM.13.0990), *L. mexicana* (LmxMNT3; LmxM.13.1210), *L. major* (LmjFNT3; LmjF.13.1210), *Trypanosoma cruzi* (TcCLBELNT3-like; TcCLB.511051.30), and *Trypanosoma brucei* subspecies, *T. b. gambiense* (TbgNT8.1; Tbg972.11.4110) and *T. b. brucei* (TbNT8.1; TB927.11.3610). MSA was generated with the T-Coffee web server using default parameters (29). Loop region, as predicted by mFold web server (21), is indicated with a box. B) Wildtype and reporter alleles at the endogenous *LdNT3* locus. Cell lines express *Fluc-BSD* under the control of either native *LdNT3* UTRs or that of a 3’-UTR harboring a scrambled stem-loop. C) Normalized Fluc activity after 24 hours of culture in the presence or absence of purine. Bars represent the mean of assays performed with 5 independent clones in technical duplicate. Single-factor ANOVA was calculated with Excel Descriptive Statistics Toolpak: ** *P* ≤ 0.01; \*\*\**P* ≤ 0.001.

We next asked whether this RNA element was sufficient to confer regulation to an *Fluc-BSD* reporter expressed from the *LdNT4* locus, which is normally not affected by purine stress. We integrated constructs encoding a Fluc-BSD reporter flanked by either wildtype *LdNT4* UTRs or a 3’-UTR harboring the *LdNT3* stem-loop at one of two different positions (Figure 3A) that were predicted via Mfold to preserve stem-loop folding (21). As shown in Figure 3B, the effects of stem-loop insertion were position-dependent. Placing the stem-loop 242 bases into the *LdNT4* 3’-UTR led to a significant decrease in basal Fluc-BSD expression, which translated to a ∼6-fold increase in luciferase activity under purine-restricted conditions. In contrast, when inserted at position +419 the stem-loop did not confer differential expression, possibly reflecting an inability of the element to fold properly in this genetic context. Thus, the *LdNT3* stem-loop is sufficient for purine-responsive expression but is sensitive to sequence context.

**Figure 3.**
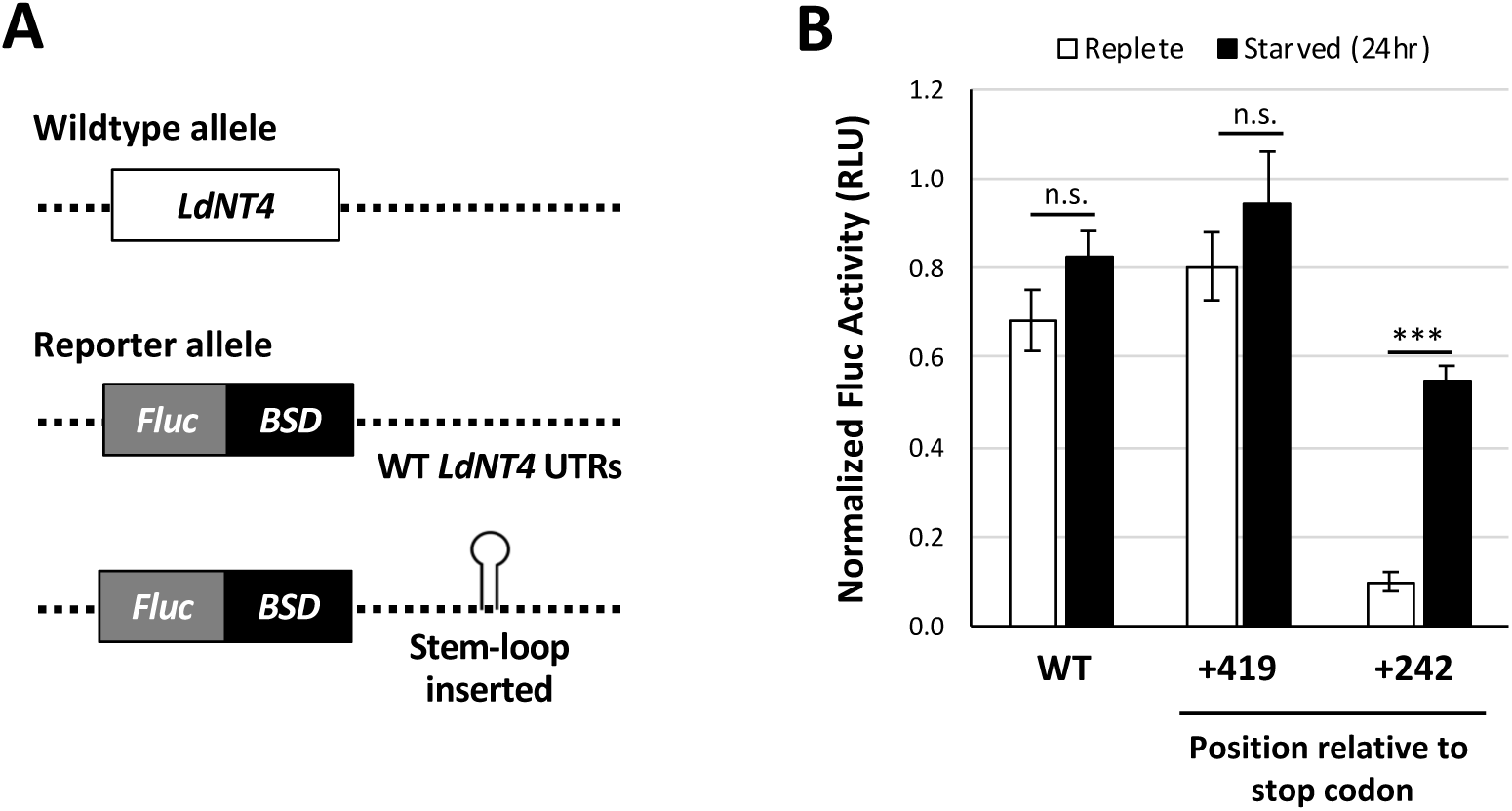
The *LdNT3* stem-loop is sufficient for purine-responsive regulation. Wildtype and reporter alleles at the endogenous *LdNT4* locus. Dashed lines indicate purine-unresponsive mRNA UTRs. In experimental cell lines, the *LdNT3* stem-loop was inserted into the *LdNT4* 3’-UTR either 242- or 419-nt downstream of the stop codon. B) Normalized Fluc activity from cell lines depicted in A, after 24 hours of culture in the presence or absence of purines. Figure shows the mean and standard deviation of experiments performed in biological and technical duplicate. Single-factor ANOVA was calculated with Excel Descriptive Statistics Toolpak: n.s., not significant, *P* ≥ 0.05; \*\*\**P* ≤ 0.001.

### Regulation by the stem-loop is species-specific and depends upon conserved residues in the loop

The purine-response elements from *TbNT8.1* and *LdNT3* share a 33 nt core with 80% identity (Figures 2A and 4A). To determine if the orthologous *TbNT8.1* stem-loop was functional in *L. donovani*, we modified the reporter construct depicted in Figure 3A to insert the minimal *TbNT8.1* stem-loop at position +242 in the *LdNT4* 3’-UTR. While this element was sufficient for regulation in *T. brucei* (16), it was unable to confer purine-responsive expression to the *Fluc-BSD* reporter in *L. donovani* (Figure 4B), suggesting that the RBPs that associate with these elements in *L. donovani* and *T. brucei* have different binding specificities.

**Figure 4.**
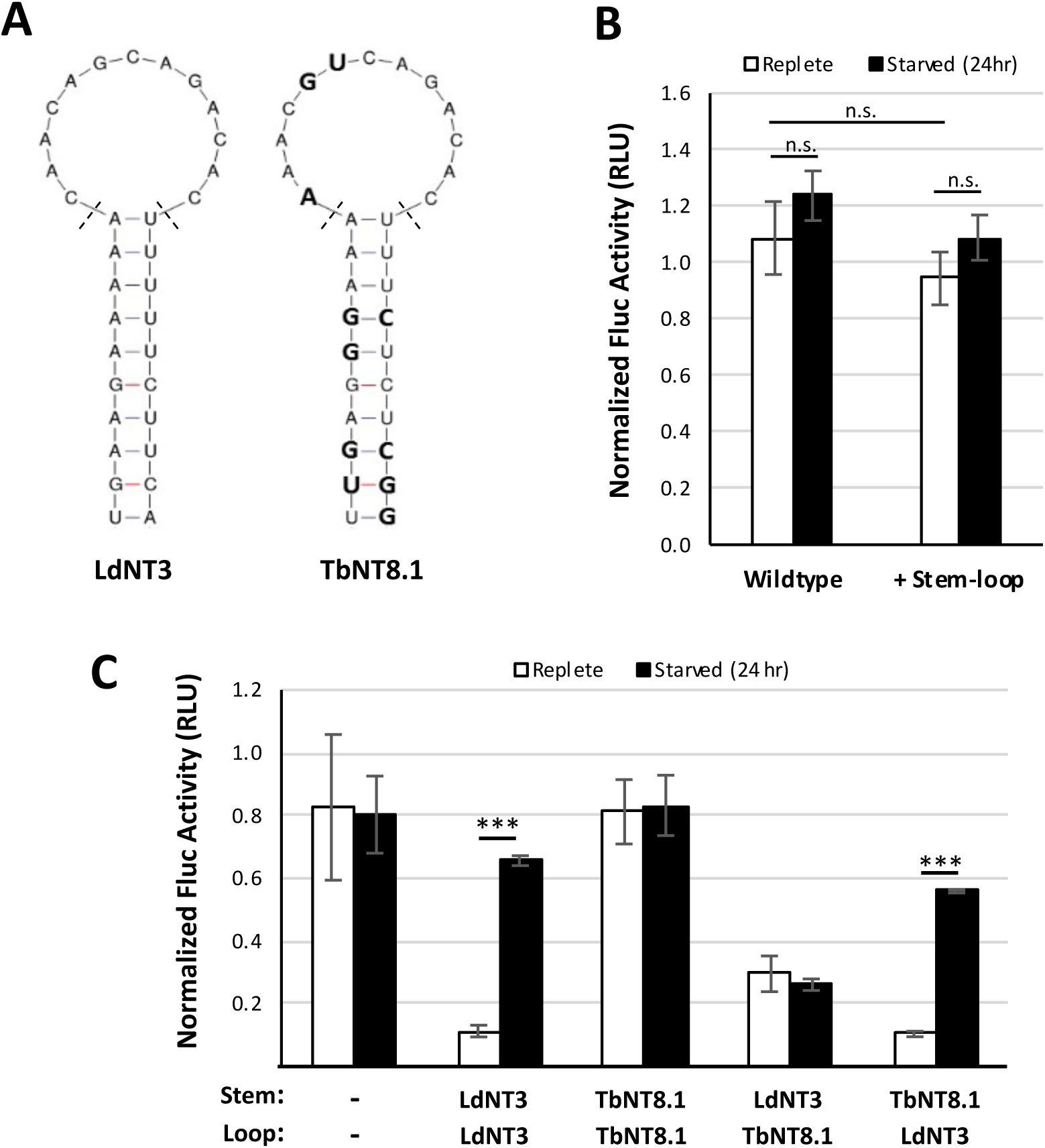
The sequence of the *LdNT3* loop, but not stem, is required for its function as a purine-response element in *L. donovani*. A) Secondary structures of the *LdNT3* and *TbNT8.1* stem-loops as predicted by mFold (21). Variant positions are highlighted on the *TbNT8.1* diagram in bold. Dashed lines delineate stem vs loop regions. B and C) Regulation by the wildtype *TbNT8.1* stem-loop (B) and stem-loop mutants (C) in *L. donovani* was tested by insertion into the *LdNT4* 3’ UTR at position +242, as depicted in Figure 3A. Bars represent the mean and standard deviation of experiments performed in biological and technical duplicate. Single-factor ANOVA was calculated with Excel Descriptive Statistics Toolpak: n.s., not significant, *P* ≥ 0.05; \*\*\**P* ≤ 0.001.

To determine if the inability of the *TbNT8.1* element to function in *Leishmania* is due to differences in the sequence of the stem, loop, or both regions, we generated chimeras in which the *TbNT8.1* loop sequence was appended to the stem of the *LdNT3* ortholog, and vice versa. These chimeric stem-loops were inserted at position +242 in the *LdNT4 Fluc-BSD* constructs and integrated into the *LdNT4* locus of a dual-luciferase compatible cell line. Parasites encoding a *TbNT8.1* loop on an *LdNT3* stem demonstrated a partial reduction in luciferase activity under purine replete conditions compared to the *TbNT8.1* stem-loop control, but there was no increase in expression upon purine starvation (Figure 4C). In contrast, Fluc-BSD expression was significantly repressed by the reciprocal *LdNT3* loop-*TbNT8.1* stem mutant in the presence of exogenous purine. This effect was reversed by 24 hours of purine stress, resulting in a similar level of luciferase induction to that conferred by the wildtype element (5.4- and 5.9-fold, respectively). These data suggest that the sequence of the *LdNT3* loop, but not stem, is essential for purine-responsive repressor activity in *L. donovani*.

We noted two distinct blocks of conservation within the loop regions of *LdNT3* stem-loop orthologs (labeled A and B in Figure 5A). We generated *LdNT3* stem-loop variants in which blocks A and B were mutated independently and tested their activity at the *LdNT4* locus. Disruption of either block resulted in a complete loss of regulation (Figure 5B), indicating that these conserved regions are important for the repressor function of the *LdNT3* stem-loop.

**Figure 5.**
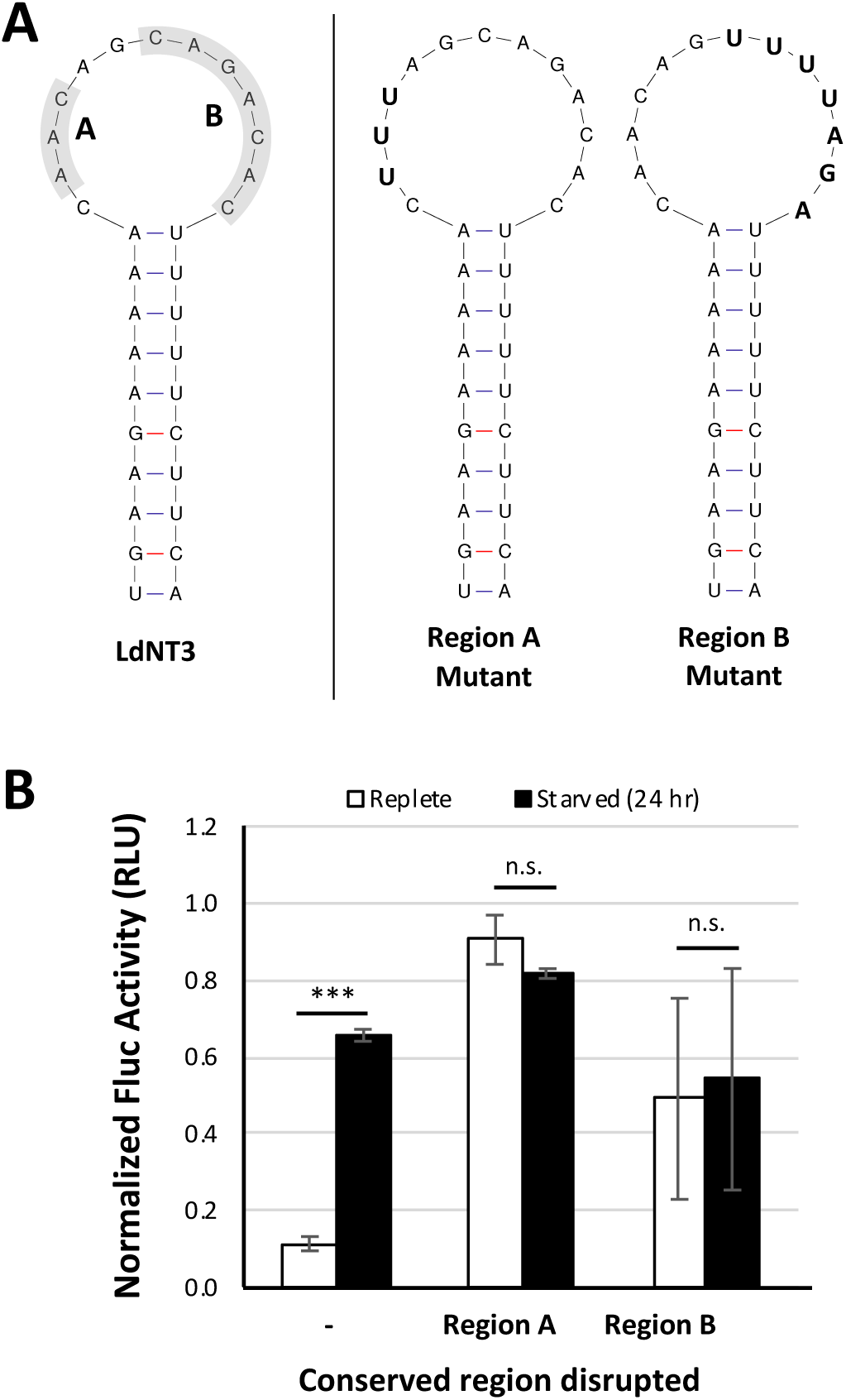
Evolutionarily conserved residues within the *LdNT3* loop are functionally important for purine-responsive gene expression. A) Evolutionarily conserved residues (Regions A and B) are highlighted in grey and their corresponding mutants are shown in boldface type. B) Regulation by Region A and B mutants was tested at the endogenous *LdNT4* locus as depicted in 3A. Bars represent the mean and standard deviation of experiments performed in biological and technical duplicate. Single-factor ANOVA was calculated with Excel Descriptive Statistics Toolpak: n.s., not significant, *P* ≥ 0.05; \*\*\**P* ≤ 0.001.

The *LdNT3* loop differs from that of *TbNT8.1* at just three positions (Figure 6A). To determine if the inactivity of the *TbNT8.1* stem-loop in *L. donovani* could be attributed to any one of these divergent bases, we generated a series of *TbNT8.1* stem-loop mutants in which each variant position was changed to the corresponding base from the leishmanial ortholog. A *TbNT8.1* stem with a wildtype *LdNT3* loop (labeled as 1-3 in Figure 6B) served as a positive control for purine-responsive induction. Like the wildtype *TbNT8.1* element, replacement of each of the three variant residues alone had no effect on luciferase activity, suggesting that species restriction is defined by multiple bases in the loop rather than any one individually. Similarly, simultaneous conversion of positions 1 and 2 to the corresponding *LdNT3* bases failed to restore purine-responsive regulation (1,2 in Fig. 6B). Two paired-position mutants (1,3 and 2,3) conferred varying degrees of repression that translated to a respective ∼2.1-fold and ∼3.7-fold increase in luciferase activity under purine stress; however, neither fully recapitulated the robust ∼7.7-fold induction observed in control cells where all three bases were changed to their *LdNT3* counterparts. Thus, each of the three nonconserved bases is important for species specificity of the repressor stem-loops, and full repressor function in *L. donovani* depends on the sequence at all three positions. Interestingly, the sequence of the 2,3 loop variant (Figure 6) corresponds to that from the orthologous *T. cruzi* transporter (Figure 2A), suggesting that the binding specificities of the cognate RNA binding proteins that associate with these stem-loops in *L. donovani* and *T. cruzi* has likely also diverged.

**Figure 6.**
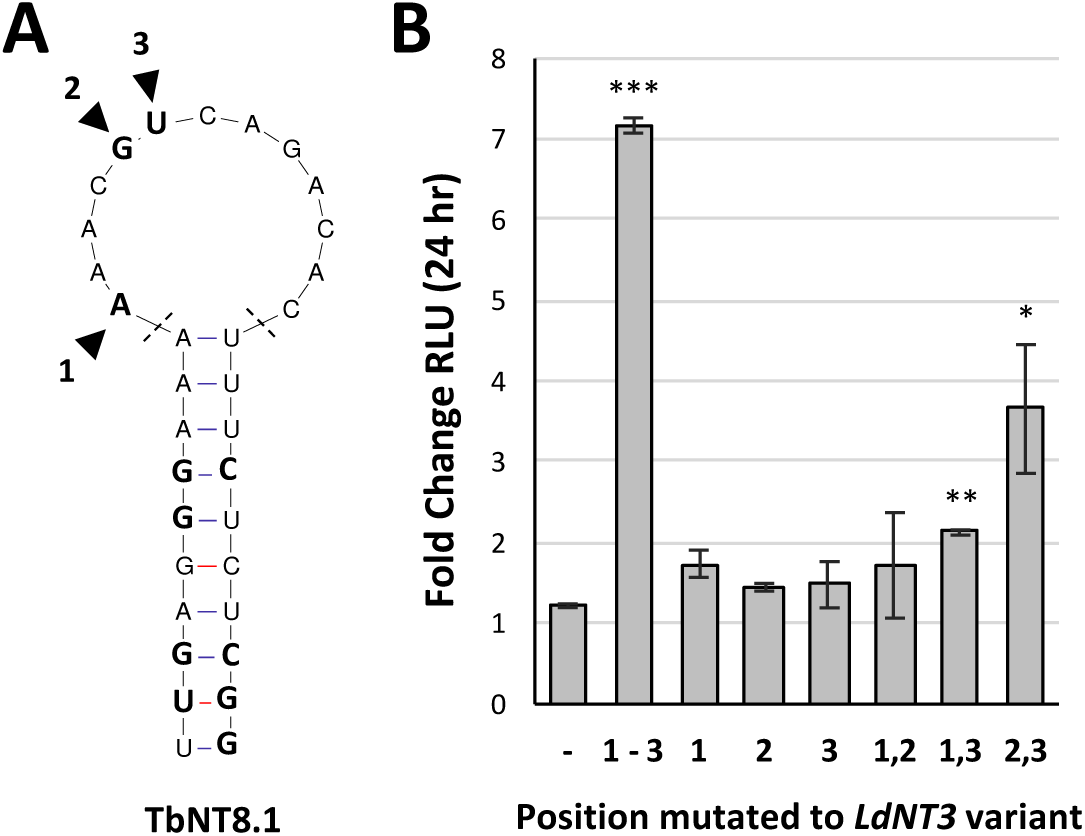
Species specificity of the purine-response element is defined by three bases within the loop. A) Three bases differentiate the *LdNT3* vs *TbNT8.1* loop. B) Regulation by single and paired-position *TbNT8.1* stem-loop mutants. Bars represent the mean and standard deviation of experiments performed in biological and technical duplicate. Asterisks (*) indicate significance. Single-factor ANOVA was calculated using Excel Descriptive Statistics Toolpak and post-test Bonferroni-corrected *P-*values are as follows: \**P* ≥ 0.007143; \*\**P* ≤ 0.001428; \*\*\**P* ≤ 0.0001428.

### Regulation conferred by the *LdNT3* stem-loop is likely mediated by a highly abundant *trans*acting factor

The steady-state mRNA level of the *Fluc-BSD* transgene flanked by *LdNT4* UTRs and harboring an *LdNT3* stem-loop is approximately 75% lower than that of the same transgene expressed from the endogenous *LdNT3* locus (Figure 7B). Having demonstrated that the *LdNT3* stem-loop is sufficient to confer purine-responsiveness to this lower-copy message, we next asked if the sequence could also mediate regulation of more abundant transcripts. The pRP vectors are a set of integrating rRNA promoter vectors generated in our laboratory that offer a range of incrementally different expression profiles in *L. donovani*. In all configurations, RNA polymerase I (Pol I) drives robust transgene transcription from the rRNA array. Graded expression is achieved using different combinations of UTRs that vary in their respective abilities to either promote or attenuate mRNA processing and stability (22). As depicted in Figure 7A, we introduced the *LdNT3* stem-loop into pRP-L_A_ and pRP-VH, low- and high-expressing vectors from the pRP suite. These constructs differ in the sequences of their 5’-UTRs but share an identical 3’-UTR derived from the *L. major* α-tubulin (*LmTUB*) intergenic region, facilitating the comparative study of the *LdNT3* stem-loop at two steady-state transcript levels while eliminating the confounding variable of local genetic context. A *Fluc* reporter gene was inserted into the multiple cloning sites of these vectors and linearized constructs were integrated into a dual-luciferase compatible cell line. As expected, the relative abundance of the *Fluc* transcript was substantially higher in these cells than when expressed from the endogenous *LdNT3* locus, to over 90- and 300-fold in the case of stem-loop-modified pRP-L_A_ and pRP-VH, respectively. Relative to control cell lines harboring an unmodified 3’ UTR, the presence of a stem-loop in either vector substantially reduced *Fluc* mRNA levels, consistent with our previous observation that this sequence negatively affects mRNA stability (Figure 7B). To test whether the *LdNT3* stem-loop could mediate purine-responsive regulation of these higher-copy transcripts, cells were cultured for 48 hours in the presence or absence of purines and subjected to dual-luciferase analysis. Fluc activity was substantially reduced in both pRP-L_A_ and pRP-VH control cells by 48 hours of purine stress, likely reflecting a general decrease in Pol I-mediated transcription of the rRNA array (Figure 7C, left). This is supported by our previous observation that Pol I protein levels are significantly downregulated in purine-starved *L. donovani* (16). Interestingly, insertion of the *LdNT3* stem-loop into the 3’-UTR of pRP-L_A_, but not pRP-VH, resulted in a moderate upregulation of luciferase activity upon starvation (Figure 7C, right), which overcame the general starvation-induced reduction in expression from the pRP vectors. The fact that the *LdNT3* stem-loop conferred purine responsive regulation to an mRNA ∼90-fold more abundant than *LdNT3* suggests that the *trans*-acting factors that associate with this element are in considerable excess of what is required solely for *LdNT3* regulation. Given that the two vectors encode identical 3’-UTRs with the *LdNT3* stem-loop inserted at the same position, the failure of the stem-loop to confer upregulation in the context of the pRP-VH construct is consistent with the possibility that the high level of mRNA expressed from this construct exceeded the availability of the cognate RNA binding protein.

**Figure 7.**
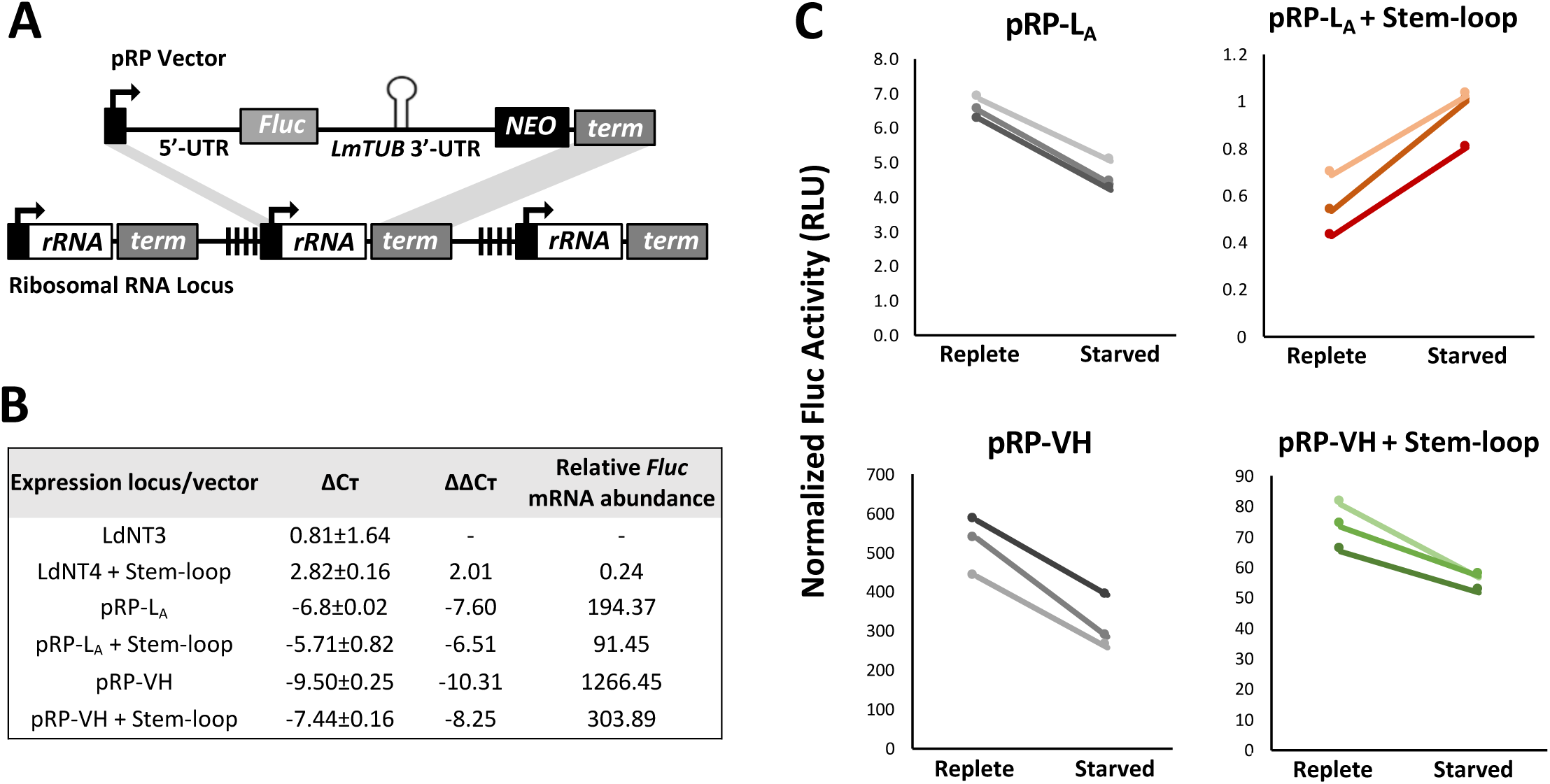
Regulation via the *LdNT3* stem-loop is likely mediated by a highly abundant *trans-*acting factor. A) Differential high-level Fluc expression achieved using integrating rRNA promoter vectors. 5’-UTRs are derived from the upstream IGRs of the *T. brucei* procyclic acidic repetitive protein and *Crithidia fasciculata* phosphoglycerate kinase B gene in pRP-L_A_ and pRP-VH, respectively. Abbreviations and symbols: term = putative rRNA terminator sequence; NEO = neomycin resistance gene; black boxes = rRNA promoter; vertical hash marks = 64 base repeats. Black arrows indicate the direction of transcription by RNA polymerase I. Grey shading indicates homology for targeted integration. B) Relative mRNA abundance for *Fluc* expressed from either the endogenous *LdNT4* locus or the rRNA locus via pRP-L_A_ and pRP-VH vectors was determined by RTq-PCR. All data were normalized to the endogenous control *UMPS*. The mean and standard deviation from two biological replicates is shown for each analysis. Transcript level is represented relative to *Fluc* expression from the endogenous *LdNT3* locus. C) Fluc activity from the pRP-L_A_ and pRP-VH vectors after 48 hours of culture in the presence or absence of purines. End points represent the mean of assays performed in technical duplicate for three independent clones of each cell line.

## DISCUSSION

Responding to the nutritional environment of the host is critical for successful parasitism. Our previous studies demonstrated that purine starvation invokes a robust nutrient stress response in *Leishmania donovani* that is characterized by a marked remodeling of the cellular proteome. We established that purine nucleoside and nucleobase transporters, including LdNT3, were highly upregulated by purine stress, and that this regulation was mediated, at least in part, via regulatory elements encoded in 5’- and/or 3’-UTRs of their mRNAs (14, 16, 18). In the current work, we have built on these observations by performing a detailed examination of the post-transcriptional and post-translational regulation of the LdNT3 purine nucleobase transporter in response to purine starvation.

Based on homology to a known purine-response element from *T. brucei* (13), we identified a 33 nt stem-loop in the 3’-UTR of *LdNT3* that serves to repress expression when purines are abundant. This is the first report of a defined nutrient stress-response element in *Leishmania.* Deletion of this element almost entirely ablated purine-responsive control by the *LdNT3* UTRs. Whereas previous work established that both *LdNT3* mRNA stability and translation are increased by purine starvation (16, 18), we found that post-translational stability of LdNT3 is not affected by extracellular purine level. Thus, in the absence of a post-translational contribution, this singular element appears to be the primary factor mediating purine-responsive changes in LdNT3 abundance.

The putative stem-loop described in this work is conserved across a variety of dixenous and monoxenous kinetoplastids including multiple *Leishmania* species, *T. brucei* subspecies, and *Trypanosoma cruzi* (highlighted in Figure 2A) as well as *Crithidia fasciculata, Leptomonas pyrrhocoris*, and *Blechomonas ayalai* (Figure S1). Conservation is particularly striking given that the relative position of the stem-loop varies substantially with respect to the stop codon and the 3’-UTRs of the orthologous transporters are otherwise poorly conserved. Thus, despite substantial expansion or contraction of the UTRs, this sequence has been maintained throughout evolutionary history, suggesting a strong selective pressure to maintain purine-responsive regulation. Interestingly, the stem-loop is absent from the orthologous purine transporter gene of the free-living kinetoplastid *Bodo saltans*, which also lacks the capacity for *de novo* purine synthesis (23). This observation may indicate that a robust purine stress response is uniquely important to the parasitic lifestyle.

As was shown for the *TbNT8.1* stem-loop in *T. brucei* (13), the *LdNT3* stem-loop is sufficient to confer regulation to an otherwise purine-unresponsive reporter in *L. donovani*. However, sequence context appears to be important, since only one of two positions into which the *LdNT3* stem-loop was inserted supported repressor activity (Fig. 3). This likely reflects differences in the propensities of the sequences surrounding the insertion points to negatively impact folding and/or accessibility of the stem-loop. The sequences of the *TbNT8.1* and *LdNT3* stem-loops differ by only 20%. We were therefore surprised to find that the two elements are not functionally equivalent. Species-restriction was attributed to just three variant bases encoded within the loop. One plausible explanation for this restriction may be that that orthologous RBPs that bind to these repressor elements have different specificities. However, as the relevant *trans*-acting factor in *T. brucei* was not identified and we have yet to isolate a candidate RBP for the *LdNT3* stem-loop, this hypothesis has yet to be directly tested.

Numerous genes are differentially expressed in response to purine stress, yet we were unable to identify the *LdNT3* purine-response element elsewhere in the *L. donovani* genome via bioinformatic analysis.^2^ However, this does not preclude the possibility that *LdNT3* is part of a purine-responsive regulon consisting of multiple genes under the control of a common RBP, as the ability of individual RBPs to interact with several disparate binding sites is well-documented (24, 25). Indeed, our observation that the stem-loop is sufficient to confer purine responsiveness to a transcript that is over 90-fold more abundant than *LdNT3* strongly suggests that its binding partner is present in substantial excess of what is required for *LdNT3* regulation, and may therefore play a role in regulating other genes within or outside of the purine stress response pathway. Despite this possibility, we suspect that the *L. donovani* purine stress response is likely mediated by multiple independent but intersecting pathways. For instance, each of the membrane purine transporters appears to be regulated by a unique combination of post-transcriptional regulatory mechanisms; the *L. donovani* purine nucleoside transporter 2 (LdNT2) is regulated solely at the translational level, while both mRNA abundance and translation are altered for purine nucleoside transporter 1 (LdNT1.1) and LdNT3 during purine stress (16). Moreover, we recently obtained evidence that the 3’-UTRs of LdNT1.1 and LdNT2 encode activator elements which, unlike the repressive stem-loop described here, promote expression when extracellular purines are depleted.^3^ Future efforts to identify the proteins that associate with these *cis*-acting elements will help to unravel the complexities of the purine stress response pathway and may uncover novel targets for therapeutic intervention.

## EXPERIMENTAL PROCEDURES

### L. donovani culture

All cell lines described herein were generated from the *L. donovani* 1S-2D clonal subline LdBob, originally obtained from Dr. Stephen Bevereley (26). LdBob promastigotes were routinely maintained at 26 °C in 5% CO_2_ and cultured in Dulbecco’s Modified Eagle-Leishmania (DME-L) medium supplemented with 5% SerumPlus™ (SAFC BioSciences/Sigma Aldrich, St. Louis, MO; a purine-free alternative to standard FBS), 1mM L-glutamine, 1x RPMI vitamin mix, 10uM folic acid, 50 ug/ml hemin, and 100 uM hypoxanthine as a purine source. For general culture maintenance, blasticidin and puromycin were used at 30 ug/ml and 25 ug/ml, respectively. To elicit purine starvation, logarithmically growing cells were pelleted via centrifugation (5000 x g for 5 min), washed once in DME-L lacking hypoxanthine but containing all other media supplements, and resuspended at a density of 2 x 10^6^ cells/ml in either purine-replete or purine-free medium.

### Luciferase constructs and cloning

All gene targeting constructs were generated using the multi-fragment ligation approach described in (27) and depicted in Figure S2. To genetically fuse *Fluc* the *LdNT3* CDS (constructs S2B and S2C), a *BSD-2A-Fluc* transgene was provided by donor vector pCRm-coBSD-2A-Fluc.^4^ The *Fluc-BSD* reporter used for gene replacement constructs (Figure S2D) was donated by pCRm-luc2-BSD (Genbank Accession number KF035118.1). 5’- and 3’-targeting sequences for integration into either the *LdNT3* or *LdNT4* locus were PCR amplified from genomic DNA with Phusion High Fidelity DNA polymerase (New England Biosciences, Ipswitch, MA) using primers listed in Table S1. For construct assembly, all vector components were digested with SfiI (or AlwNI, where indicated), gel-purified, and combined in a single ligation step as depicted in Figure S2A.

To generate *LdNT3* stem-loop-deficient mutants, the *LdNT3-*targeting construct shown in Figure S2D was modified with the QuikChange Site-directed Mutagenesis Kit (Stratagene, Lajolla, CA) using the primers listed in Table S2. To insert the *LdNT3* stem-loop and variations thereof into the *LdNT4* 3’-UTR, the *LdNT4* version of construct S2D was subjected to whole-plasmid PCR amplification via Phusion polymerase using the primers listed in Table S3. PCR products were DpnI-treated to eliminate template plasmid and circularized with NEBuilder HiFi DNA Assembly Master Mix (New England Biosciences, Ipswitch, MA) according the manufacturer’s instruction.

All primers used to modify pRP vectors are listed in Table S4. To integrate the firefly luciferase gene into the rRNA array, *Fluc* was amplified from pCRm-luc2-BSD and cloned into the SfiI sites of pRP-L_A_ and pRP-VH (22). Insertion of the *LdNT3* stem-loop into pRP-L_A_ and pRP-VH was accomplished in a step-wise fashion. First, the vectors were subjected to whole-plasmid amplification to introduce two nonidentical BstXI sites into their shared 3’-UTRs (see primer sequences for detail). A version of the *LdNT3* stem-loop was synthesized with flanking BstXI and PCR primer binding sites (Genscript, Piscataway, NJ) and inserted into the BstXI sites of the modified pRP vector.

### Transfections

To generate dual-luciferase cell lines, an LdBob derivative expressing *Rluc* from the endogenous *UMPS* locus was used as a recipient for all vector transfections (18). Transfections of mid-log stage promastigotes were performed with ∼3ug SwaI-linearized plasmid DNA using the high-voltage electroporation protocol described by Robinson and Beverley (28). Immediately following electroporation, cells were transferred into 5 mL of complete DME-L and 200 ul was added to the first column of wells on a 96 well plate and subjected to 2-fold serial dilution to derive independent clones. Transfections were incubated overnight at 26 °C in 5% CO_2_. Selection was initiated the following day by adding 100 ul of 2X blasticidin (60 ug/ml) or, for integration of pRP-L_A_ and pRP-VH vectors harboring *NEO*, G418 (50 ug/ml) to each well. Proper integration of the constructs was verified via PCR for all clones.

### Dual-Luciferase analysis

Firefly and *Renilla* luciferase activities were assessed using the Dual-Glo Luciferase Assay System from Promega. Analyses were performed using 35 ul of cell culture in white polystyrene 96-well half-area plates (Corning, Amsterdam) as described in the product technical manual. For each incubation step, plates were protected from light and shaken for 10 minutes at room temperature on an orbital shaker. Luminescence was measured using a Veritas Microplate Luminometer (Turner BioSystems, Sunnyvale, CA).

### RTq-PCR Analyses

Total cellular RNA was isolated from 5 × 10^6^ log-stage parasites using the Qiagen RNeasy Plus Micro Kit following the protocol for animal and human cells. Cell lysates were disrupted using a QIAshredder spin column. To eliminate contaminating genomic DNA, RNA samples were subjected to DNaseI digestion using the TURBO DNA-*free* kit. First-strand cDNA synthesis was performed using 1ug of RNA template with the High Capacity cDNA Reverse Transcription Kit. cDNA samples were subsequently diluted so as to reduce the total input RNA to 6 ng per ul. Dye-based real-time qPCR was performed with NEB Luna Universal qPCR Master Mix according to the manufacturer’s instructions using 2 ul (12 ng) of diluted cDNA. Previously validated PCR primers (16) are listed in Table S5. Reactions were run on an Applied Biosystems StepOnePlus instrument using the “Fast” ramp speed and the following thermocycling parameters: 95 °C for 60 seconds; 40 cycles of denaturing at 95 °C for 15 seconds followed by a 30 second extension at 60 °C. A final melt curve step was included to verify the specificity of amplification. The relative abundance of the *Fluc* transcript expressed from various genetic loci was determined using the comparative CT (ΔΔCT) method as described (16).

## Acknowledgements

We thank Drs. Scott Landfear and Georgiana Purdy for helpful comments on the manuscript. The content is solely the responsibility of the authors and does not necessarily represent the official views of the National Institutes of Health.

## *In order of appearance, the abbreviations used are

CDS: coding sequence;
RBP: RNA-binding protein;
LdNT3: *L. donovani* purine nucleobase transporter 3;
nt: nucleotide;
Fluc: firefly luciferase;
IGR: intergenic region;
Rluc: *Renilla* luciferase;
UMPS: UMP synthase;
BSD: blasticidin resistance gene;
2A: *Thosea asigna* virus 2A peptide;
LdNT4: *L. donovani* purine nucleobase transporter4;
TbNT8.1: *T. brucei* purine nucleobase transporter 8.1;
Pol I: RNA polymerase I;
LmTUB: *L. major* α-tubulin;
DME-L: Dulbecco’s Modified Eagle-Leishmania medium;
RT-qPCR: Reverse transcription quantitative PCR;
PAC: puromycin resistance gene;
term: rRNA terminator sequence;
NEO: neomycin resistance gene

## Footnotes

1 Data not shown.

2 M. H. Licon and P. Yates, unpublished observation.

3 M. H. Licon and P. Yates, unpublished observation.

4 P. Yates, Manuscript in preparation

